# Tirzepatide inhibits tumor growth in mice with diet-induced obesity

**DOI:** 10.1101/2023.06.22.546093

**Authors:** Linxuan Huang, Jibin Zeng, Ye Wang, Michael Pollak

## Abstract

Tirzepatide, a drug used in management of type II diabetes, is an activator of both glucose-dependent insulinotropic polypeptide (GIP) and glucagon-like peptide 1 (GLP-1) receptors. Tirzepatide treatment leads to weight loss in murine models of obesity, and clinical trials have shown the drug can lead to weight loss up to ∼ 20% in overweight patients. Obesity has been shown to increase risk and/or to worsen prognosis of certain common cancers, including colon cancer, but the effect of tirzepatide on neoplasia has not been examined in detail. We studied the effects of this drug on the murine MC38 colon cancer model, which has previously shown to exhibit accelerated growth in hosts with diet-induced obesity. Tirzepatide did not cause tumor regression, but reduced tumor growth rates by ∼ 50%. This was associated with substantial reductions in food intake, and in circulating levels of insulin and leptin. Tirzepatide had no effect on MC38 cancer cell proliferation *in vitro*, and the effect of tirzepatide on tumor growth *in vivo* could be phenocopied in placebo treated mice simply by restricting food intake to the amount consumed mice receiving the drug. This provides evidence that the drug acts indirectly to inhibit tumor growth. Our findings raise the possibility that use of tirzepatide or similar agents may benefit patients with obesity-related cancers.

## Introduction

Obesity is a leading public health challenge. Increased risk and/or worse prognosis of certain cancers are associated with obesity, and the global cancer burden attributable to obesity is substantial (1-4). The mechanisms underlying the influence of obesity on neoplastic disease are complex and incompletely characterized. There is substantial evidence that hormonal changes associated with obesity play a role. The hyperinsulinemia resulting from obesity-associated insulin resistance represents one plausible mediator, as many cancers in patients with systemic insulin resistance express insulin receptors and are insulin-sensitive as they are growth-stimulated by activation of signal transduction pathways downstream of the insulin receptor (5). Mendelian randomization studies (for example (6)) provide evidence that hyperinsulinemia (rather than hyperglycemia) plays a causal role in obesity-related neoplasia. However, it is unlikely that the obesity – neoplasia association results from any single factor and there also is evidence for roles for leptin (7) immune factors (8, 9), adipocyte-cancer interactions (10, 11), and other mechanisms.

Tirzepatide is a dual GIP and GLP-1 receptor agonist first studied clinically in 2018 (12, 13). In common with other incretin-based drugs, it is effective for treatment of type II diabetes (14-17), and leads to substantial weight loss in obese subjects with or without T2D (18). While incretins were originally characterized as enhancers of insulin secretion, the actions of incretin-related drugs are pleomorphic and include reduced appetite, reduced weight, reduced insulin resistance, and reduced insulin levels (19-21). The reduction in appetite involves hypothalamic effects (22).

Early in the development of incretin-based drugs, concerns were raised concerning the possibility that these agents might increase risk of certain cancers (23), but most studies of this issue have been reassuring (24, 25). Despite these early concerns, we hypothesized that the effects of tirzepatide on obesity and associated metabolic abnormalities would lead to anti-neoplastic activity, at least for a subset of cancers. As models of carcinogenesis and cancer risk require at least a year of observation, our initial approach was to study effects of tirzepatide on growth rate of pre-existing cancers. We used the mc38 murine colorectal cancer model, previously shown to exhibit enhanced growth in C57/B mice with diet-induced obesity (26, 27).

## Methods

### Animals

All experiments were approved by the McGill University Animal Care Committee. Male C57BL/6 mice with diet-induced obesity (DIO) were purchased from Taconic (NY, USA) and put on either a control diet (Teklad, #2918) or a high-fat diet (Research Diets, D12492) for 2-4 weeks. The control diet consisted of 18.4% protein, 6% fat, 44.2% carbohydrate. The high-fat diet consisted of 20% protein, 20% carbohydrate, and 60% fat. Animals were housed in a temperature-controlled (23-25°C) facility with a 12:12 light:dark cycle and had a one-week acclimation prior to the experiments.

### Tirzepatide

Tirzepatide was purchased from Selleckchem (#P1206, TX, USA). To formulate the solution from the peptide powder form, per 1 mg of the powder was added to 1 mL of the buffer (Tris-HCl pH8.0 + 0.02% PS80). The solution was then poured through a sterile filter (Milipore) and divided into aliquots and stored at -80 degrees.

### *In vivo* work

Animals were randomized into two groups based on their weight, and received vehicle (Tris-HCl pH8.0 + 0.02% PS80) or tirzepatide treatment by subcutaneous injection daily. The doses for the vehicle and tirzepatide groups were 10 mL/kg and 10 nmol/kg, respectively. Mouse body weight and food consumption were measured 2 times a week. In allograft experiments, prior to the tumor injection, cells were resuspended in PBS. Mice were injected subcutaneously with 2×10^5^ cells in a volume of ∼ 100 uL. Tumors were palpable in most of the mice after ∼ one week of the injection. Tumor size was measured every other day with an electronic caliper.

### Cell lines and culture conditions

MC38 cells were obtained from ATCC. Cells were cultured in flasks in 10% FBS-DMEM. For *in vitro* studies, 2×10^4^ MC38 cells per well were plated in 12-well plates with different concentrations of tirzepatide (0, 35, 70, 100 nM diluted with 10% FBS-DMEM from 1 M stock solution). Cells were cultured for 96 hour, with a media change at 48 hours.

### Hormone measurements

ELISA kits for insulin, c-peptide, and leptin were purchased from R&D systems. Blood was collected at the end of the experiment through cardiac puncture in heparinized capillary tubes. Plasma was drawn from the supernatant after centrifugation.

### Statistical analyses

Paired or unpaired Student’s *t*-test was used to compare the differences between two groups. All statistical analyses were performed using Graphpad Prism 9.0 (MA, USA).

## Results

### Tirzepatide reduces body weight and food consumption in mice

To confirm prior reports (13, 20), we examined the effect of tirzepatide on weight and food consumption. The weight-loss effects of tirzepatide were confirmed in diet-induced obese (DIO) mice (Figure 1A). The group receiving daily tirzepatide treatment exhibited a significant decrease in weight compared to the group receiving vehicle injections. Food consumption was also measured, and demonstrated lower average food intake in the tirzepatide-treated group (Figure 1B). The average daily energy intake in the control mice was 16.8 kcal while that in the tirzepatide treated mice was 11.3 kcal.

**Figure 1:**
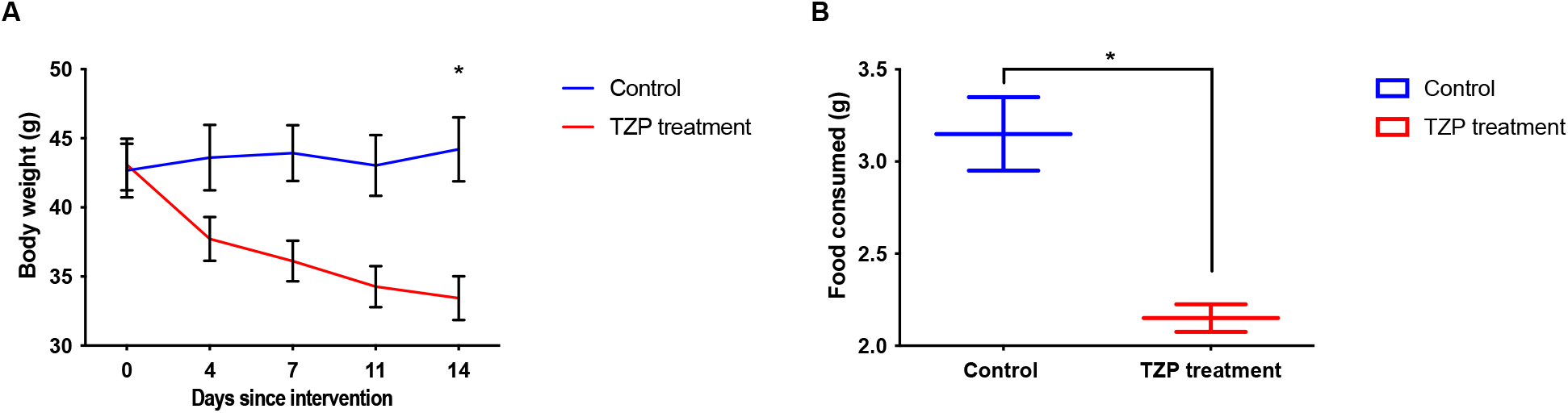
Tirzepatide reduces body weight and average daily food consumption in mice. (A) Changes in mouse weight during 14 day period on either vehicle (n = 4) or tirzepatide (n = 4). (B) Average weight of daily food consumed per mouse. Data are presented as mean ± SEM. \**P* < 0.05 compared with control.

### Animals receiving tirzepatide treatment showed lower glucose, insulin, leptin, and c-peptide levels compared to the control group in fed and fasted states

After 15 days of treatment, mice were sacrificed, and blood samples were collected for hormone analyses. ELISA results demonstrated that mice treated with tirzepatide displayed lower levels of glucose concentration, insulin, c-peptide, and leptin (Figure 2) compared to untreated controls. The decline in leptin levels may reflect a tirzepatide-induced reduction in leptin resistance (28).

**Figure 2:**
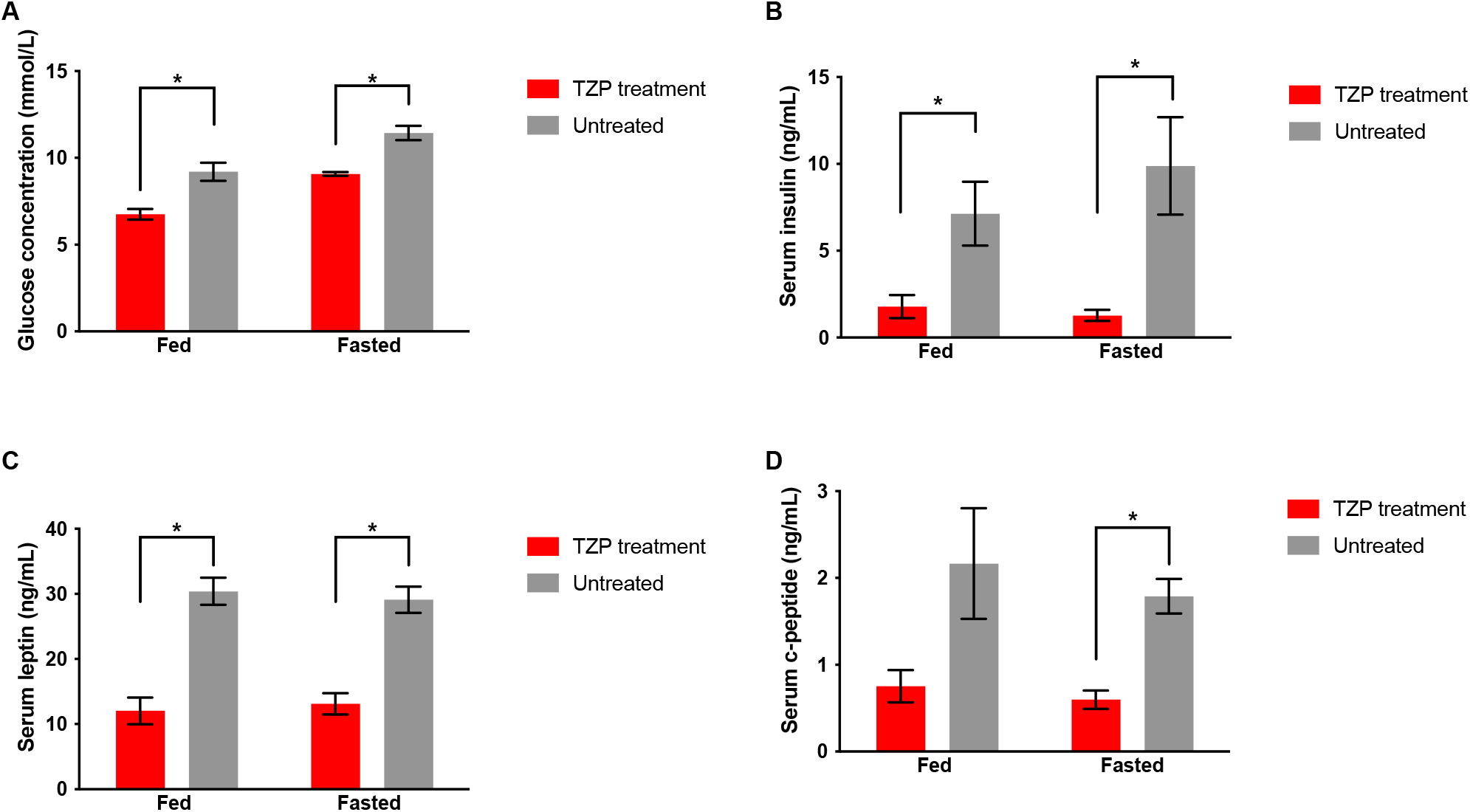
Mice receiving tirzepatide treatment have lower glucose, insulin, leptin, and c-peptide levels compared to the untreated group in fed and fasted states. (A) Blood glucose concentration, (B) serum insulin level, (C) serum leptin levels, and (D) serum c-peptide levels were measured in tirzepatide treated group (n = 4) and baseline group (n = 3) during the fed state and fasted state on the last day of the experiment (day 15). Data are presented as mean ± SEM. \**P* < 0.05 compared with control or baseline.

Patients with obesity-related insulin resistance generally exhibit elevated insulin levels compared to healthy individuals. A prior publication revealed that chronic treatment with tirzepatide reduced insulin levels in obese mice (20). To explore the relationship between body weight and serum insulin, a scatter plot analysis was conducted (Figure 3). The slope of the line relating weight to insulin concentration indicated the expected relationship for within-group variation among the controls, which associated higher weight with higher insulin levels. In contrast, mice in the tirzepatide-treated group not only had lighter body weight, but their insulin levels did not vary with their weight, as was seen in the control mice. While we detected these differences even with a small sample size, confirmatory experiments using more mice are underway.

**Figure 3:**
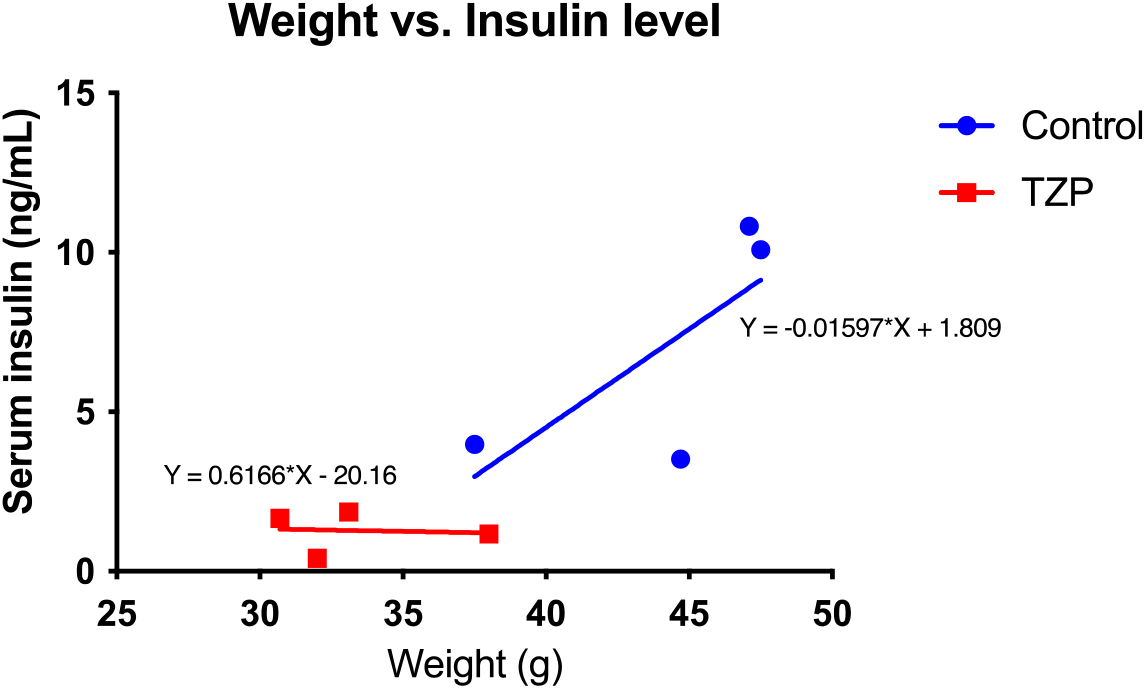
Scatter plot of insulin level vs. mouse body weight for mice receiving TZP or vehicle.

### Effect of tirzepatide or pair-feeding on MC38 tumor growth

Obese C57BL/6 mice on a high-fat diet with MC38 allografts were used in this experiment. We compared tumor growth between mice (***a***) on an *ad lib* high-fat diet given daily subcutaneous vehicle injections (controls) to that in (***b***) mice on the same diet given tirzepatide (10 nmol/kg) daily by subcutaneous injection and to (***c***) that in a pair-fed group given vehicle injections with daily food intake restricted to the amount consumed by tirzepatide-treated mice. The animals were treated according to their assigned groups for 3 days prior to the tumor injection. The animals continued to receive daily injections, and on day 14, tumors started to be palpable. 25 days after tumor injection, animals were sacrificed and blood and tumors were collected on day 28 (Figure 4A). The tirzepatide-treated group and the pair-fed group both exhibited significant weight reduction (Figure 4B). The control group had significantly larger tumor volumes compared to both the tirzepatide-treated group and the pair-fed group (Figure 4C).

**Figure 4:**
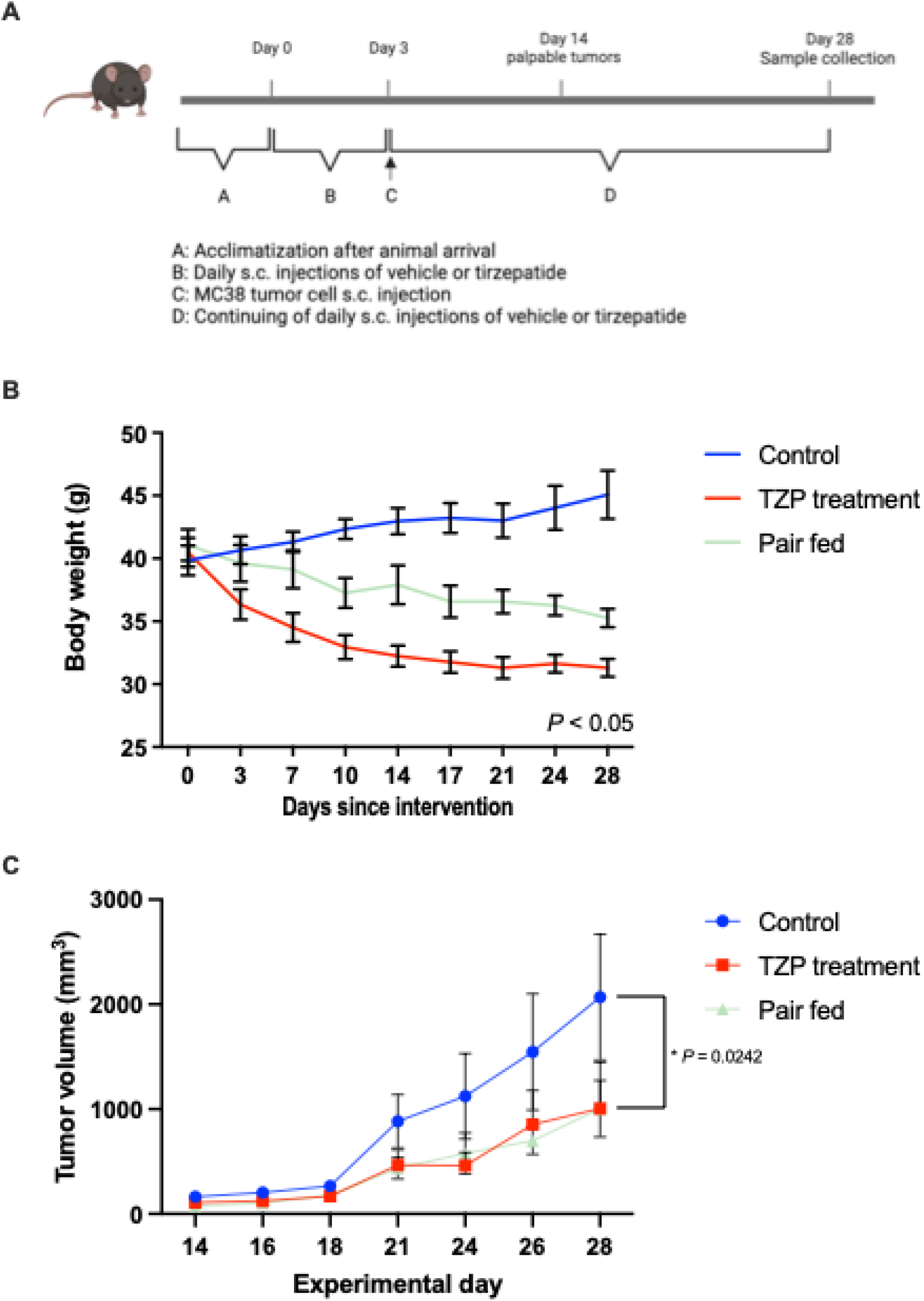
Both body weight and tumor growth are reduced in mice treated with tirzepatide or pair fed with the amount of food consumed by tirzepatide-treated mice. (A) Schematic graph of the experiment timeline. (B) Body weight change among the three groups: control (n = 6), tirzepatide treated (n = 6), and pair fed (n = 6). The pair fed group received vehicle injection but the allowed daily food consumption was limited to that consumed by the tirzepatide-treated mice. (C) Tumor volume change from experiment day 14 to day 28. Data are presented as mean ± SEM. \**P* < 0.05 compared with control or baseline.

### Tirzepatide has no direct effect on MC38 proliferation *in vitro*

Given the significant inhibition of MC38 tumor growth *in vivo* caused by tirzepatide, we investigated its impact on MC38 proliferation *in vitro* to determine if the drug acted directly as a growth inhibitor. MC38 tumor cells were cultured in different concentrations of tirzepatide for 96 hours to assess any direct effects on cell growth (Figure 5). The results indicate that tirzepatide at concentrations achieved in patients does not significantly influence MC38 cell proliferation *in vitro*.

**Figure 5:**
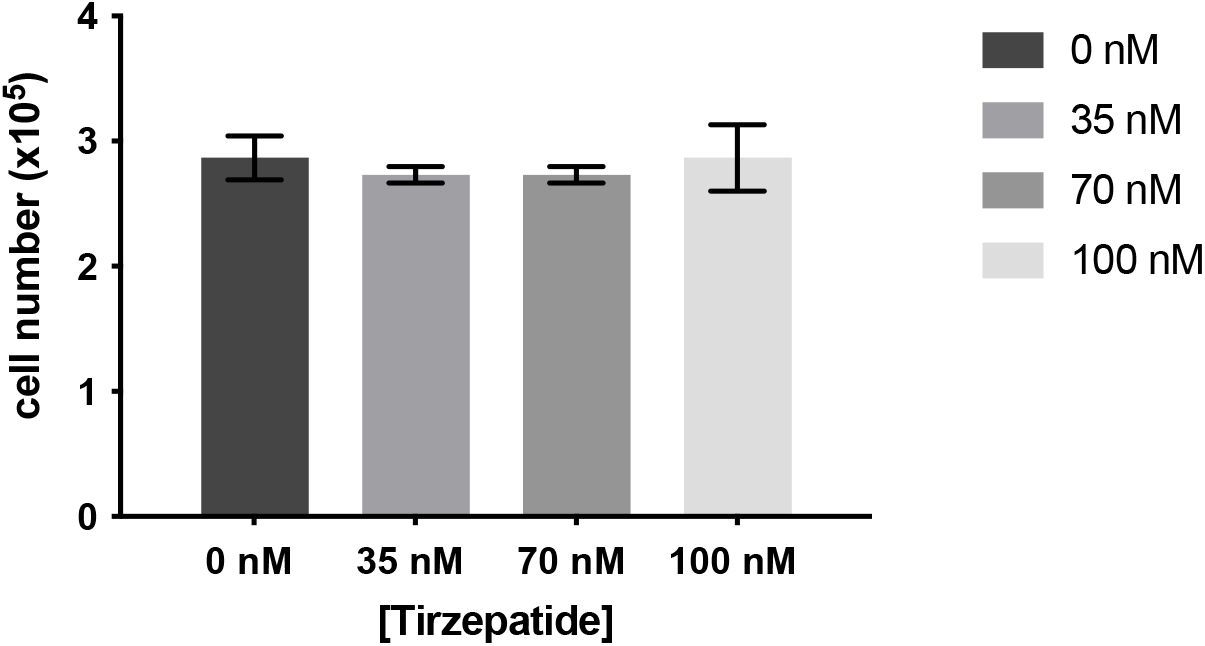
Tirzepatide does not change MC38 proliferation *in vitro* after 96 hours. Data are presented as mean ± SEM.

## Discussion

While some aggressive cancers proliferate rapidly in an automatous fashion, the behavior of others is influenced by host endocrine factors. Therapeutic exploitation of neoplastic dependency on host factors dates back to ovariectomy being used as the first systemic treatment for cancer (29), and targeting estrogen and androgen signaling remain central in the treatment of breast and prostate cancer. The paradigm of hormonal dependency of cancer may extend beyond steroid hormones to nutritionally-regulated peptide hormones such as insulin and leptin (5, 7), and may mediate at least in part the classic observation that severe reduction of food intake inhibits carcinogenesis in rodent models (for example (30)).

Notwithstanding the morbidity associated with weight loss and cachexia seen in advanced metastatic cancer, there is interest in the hypotheses that weight loss by obese individuals might reduce the risk of a future cancer diagnosis and/or that weight loss by obese people with early cancers might improve prognosis. Rigorous clinical evidence to support these hypotheses is lacking, except for the intriguing finding that cancer risk is significantly reduced following treatment of morbid obesity by bariatric surgery (for example (31). Many studies have studied the feasibility of short-term weight loss by dieting and/or exercise among patients with early cancer, but sustained weight reduction remains challenging, as does long-term compliance with interventions such as intermittent fasting or ketogenic diets. In this context, the use of incretin drugs to achieve substantial and sustained weight loss is appealing, and might make clinical trials with cancer endpoints (as distinct from short-term trials with biomarker endpoints) feasible.

Our findings confirm that tirzepatide treatment leads to reduced food intake, weight loss, and to substantial reductions in leptin and insulin levels. We also observed that MC38 tumor growth was substantially reduced in tirzepatide-treated mice as compared to vehicle treated controls. The observation that tirzepatide had no effect on *in vitro* growth of MC38 cells, together with the finding that the effect on *in vivo* tumor growth could be phenocopied simply by restricting food intake in vehicle-treated mice to that which tirzepatide -treated animals chose to eat, strongly suggests that the mechanism of action is indirect.

The reductions in insulin and leptin levels seen in tirzepatide treated animals are strong candidate mediators of the effects of the drug on tumor growth. It is noteworthy that reduction of insulin receptor activation of insulin-stimulated cancer cells has previously been proposed as a therapeutic strategy (reviewed in (5)), but small molecule insulin receptor antagonists were not clinically useful, likely because dosing had to be limited to levels that would not cause hyperglycemia (32, 33). Metformin was also extensively studied as an anti-cancer drug, based in part on its ability to reduce weight and insulin levels, but clinical trials were disappointing (34, 35), perhaps because effects on insulin levels were modest in magnitude. Ongoing studies are comparing effects of tirzepatide and other incretin-type drugs to those of metformin, SGLT2 inhibitors and caloric restriction regimes on insulin levels and insulin receptor activation in neoplastic tissue, as there is interest in determining the most effective and practical strategy to lower insulin levels for various potential indications in oncology, including increasing efficacy of PI3K inhibitors (36, 37).

Pharmacoepidemiologic evidence concerning effects of tirzepatide on cancer risk or prognosis are lacking because available follow-up periods are too short. Other incretin drugs have been found in some, but not all, pharmacoepidemiologic studies to lead to reduced risk of cancer (25). Our experimental results provide additional evidence that justifies clinical and population-based studies of the influence of incretin drugs on neoplastic disease in obese patients. Future studies will include confirmation in additional models of obesity-stimulated cancer growth, and analysis of changes in signal transduction networks within tumor allografts following tirzepatide treatment, in order to directly address our hypothesis of an indirect mechanism of action.

## Notes

### Competing Interest Statement

The authors have declared no competing interest.

